# The use of ‘artificial saliva’ as a neutral control condition in gustatory research

**DOI:** 10.1101/2019.12.18.880880

**Authors:** Imca S. Hensels, Deborah Talmi, Stephanie Baines

## Abstract

Distilled water with NaHCO_3_ and KCl, a solution often referred to as ‘artificial saliva’ because its chemical composition mimics human saliva, is the control liquid of choice in gustatory research, because it is believed to be entirely affectively neutral. Yet evidence that human research volunteers perceive this liquid as emotionally neutral is lacking. Unpublished data from our lab suggested that this solution might be perceived as aversive. This study set out to systematically test the parameters influencing taste neutrality. We used two different concentrations of distilled water with NaHCO_3_ and KCl, as well as bottled water as a control taste. Healthy adults rated all tastes on two separate scales to rule out an interpretation based on the specifics of a single scale. Our participants rated artificial saliva as aversive on both scales. The bottled water was rated as neutral on both scales, and as significantly less intense in sensation than both concentrations of the distilled water solution. This is the first study to have directly tested the subjective feelings that accompany the ingestion of these oft-used solutions. We found that these liquids, which were previously assumed to be neutral, may not be perceived as such by research participants. Therefore, future gustatory studies should take care when using this solution as a neutral baseline. It is advised that trial-by-trial ratings are collected. Also, depending on the nature of future studies, bottled water may be considered as a preferable neutral baseline.

## 1. Introduction

Most experiments of gustatory function, as well as those that employ gustatory stimuli to manipulate affective and cognitive function, need a neutral baseline. A prevalent neutral baseline is distilled water with KCl and NaHCO_3_ (see Franken et al., 2011; Grabenhorst, D’Souza, Parris, Rolls, & Passingham, 2010; Grabenhorst et al., 2008; Hird et al., 2017; Nakamura et al., 2011; Rolls, Kellerhals, & Nichols, 2015). This compound is typically referred to as ‘artificial saliva’ because its chemical composition mimics that of human saliva. Whereas humans can sense water in their mouths, distilled water with NaHCO_3_ and KCl is designed to not be sensed, and therefore to be less sensorily stimulating than normal water to human research volunteers. These properties led to its frequent use as a chemosensorily neutral baseline in research that employs gustatory stimuli (Franken et al., 2011).

However, it is unclear whether ‘artificial saliva’ is indeed perceived as neutral by research participants, as no previous studies have reported collecting ratings of the pleasantness or intensity of its perceived taste. The aim of the present study was, therefore, to test the pleasantness and intensity of this liquid in order to establish whether distilled water with NaHCO_3_ and KCl is indeed considered to be emotionally neutral. This is important because if it is, then this would validate the use of this liquid in the literature. By contrast, if the current study shows that this liquid is perceived to be pleasant or unpleasant then this would have consequences for its use as a neutral control stimulus in future research.

One important factor that needs to be taken into account when collecting gustatory pleasantness ratings is the rating scale that is used. This is because it has been found that the size of measured differences in taste perception between different flavours depends on the scale that is used (Kalva et al., 2014). In this study, we therefore compared two rating scales that are used frequently in gustatory research to ensure that scale choice did not introduce systematic bias in ratings. We used a nine-point rating scale, ranging from one being not pleasant at all, to nine being very pleasant (Kalva, Sims, Puentes, Snyder, & Bartoshuk, 2014). We included this scale for because of its ease of use – a simple nine-point scale on which to rate each liquid. The second scale we used was the hedonic general labelled magnitude scale (hedonic gLMS; Bartoshuk, Catalanotto, Hoffman, Logan, & Snyder, 2012), which is a logarithmic scale. This scale is the most prevalent in gustatory research.

Gustatory research utilises two different concentrations of distilled water with NaHCO_3_ and KCL to make ‘artificial saliva’: 2 mM NaHCO_3_/15 mM KCl or 2.5 mM NaHCO_3_/ 25 mM KCl. We tested both of these concentrations here to determine whether the hedonic perception of the liquid depends on the strength of the concentration. Both concentrations were also compared to bottled water as an alternative neutral taste. Bottled water is not as chemically similar to saliva as distilled water with added KCl and NaHCO_3_. Therefore, it is thought to be sensed more intensely in the mouth (Franken et al., 2011). However, bottled water is a frequently-consumed liquid that intuitively one might consider hedonically ‘neutral’ in the common sense of the word. Indeed, the familiarity of bottled water might be an influential factor in the subjective experience of neutrality. Given that bottled water is a common liquid that we expected the participants to already be familiar with, this liquid was also used by participants to rinse their mouths with between trials. For this study, we hypothesised that the distilled water with differing concentrations of KCl and NaHCO_3_ would all be rated as more unpleasant and more intense than bottled water.

## 2. Methods

### 2.1 Participants

30 participants (of whom seven were males) were recruited for this study. Participants could take part if they were age 18 or over, non-smokers, with no history of neurological or psychiatric illness (except for binge eating disorder), not currently on a restrictive diet, and with no food allergies that would interfere with the stimuli used in this study. Three participants were excluded from the final data analysis because one or more of their neutral valence ratings were more than three standard deviations away from the mean, leaving 27 participants in the final sample. Descriptive statistics for the sample are given in Table 1.

### 2.2 Materials

#### 2.2.1 Stimuli

Three types of nominally-neutral liquids were used. Two of these are the two most commonly-used control liquids in gustatory research: distilled water with either 2 mM NaHCO_3_ and 15 mM KCl or with 2.5 mM NaHCO_3_ and 25 mM KCl (Franken et al., 2011; Hird et al., 2017; Bloemendaal et al., 2015; Bohon et al., 2009; Grabenhorst, Rolls, Parris, & D’Souza, 2010; Iannilli, Noennig, Hummel, & Schoenfeld, 2014; Kringelbach, O’Doherty, Rolls, & Andrews, 2003; McCabe & Rolls, 2007; Nakamura et al., 2011; O’Doherty, Deichmann, Critchley, & Dolan, 2002; Rolls et al., 2015; Rolls, 2009; Stice, Burger, & Yokum, 2013; Stice et al., 2008; Sun et al., 2014; Wang, Yang, Hajnal, & Rogers, 2015). The third was unflavoured, Tesco own-brand, still bottled water. These three nominally-neutral liquids were compared to a pleasant and an unpleasant taste to anchor both scales. Apple juice (Tesco-own brand, from concentrate) was used as the pleasant taste, while the unpleasant taste was bottled water with 5 grams of salt (NaCl) per 100ml.

#### 2.2.2 Scales

The two scales used in this study were the nine-point rating scale (1=not pleasant at all, to 9=very pleasant) and the gLMS (see Figure 1; Kalva et al., 2014). There were two versions of each scale: one to measure valence, and one to measure intensity.

**Figure 1.**
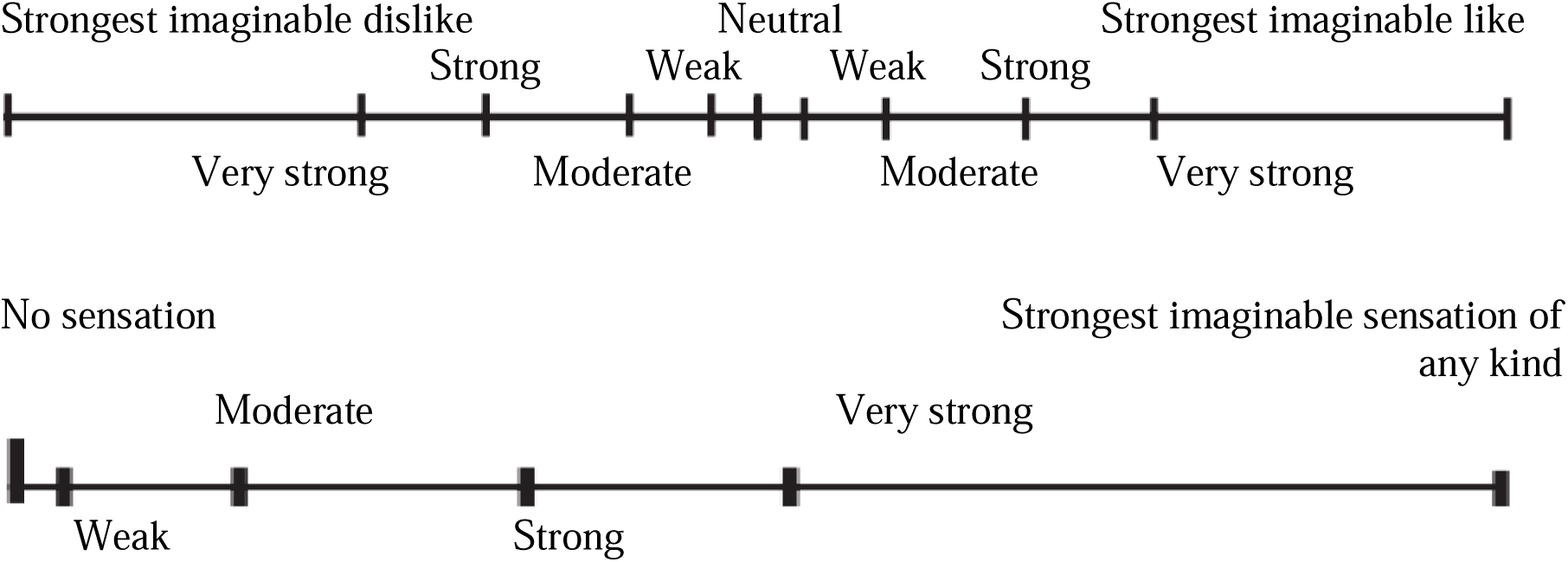
Example of the gLMS rating scale, showing the hedonic gLMS above and the sensory gLMS below.

The “hedonic nine-point scale”, measuring valence, had three anchors in this study: 1 (“not pleasant at all”), 5 (“neutral), and 9 (“very pleasant”). The “sensory nine-point scale”, measuring intensity had two anchors: 1 (“not intense at all”) and 9 (“very intense”).

The “hedonic gLMS”, measuring valence, is a continuous scale that ranges from “strongest imaginable disliking of any kind” to “strongest imaginable liking of any kind” (Kalva et al., 2014). It requires people to mark an X on the scale based on where they think the preceding stimulus falls on this continuum. Participants are first informed that the “strongest imaginable (dis)liking of any kind” can be any sensory feeling, and does not need to be related to taste. They are instructed that they should rate the taste stimuli against this feeling. This scale was scored by measuring the distance (in millimetres (mm)) between the X put on the scale by the participant and the ‘neutral’ marker. In the current study, the hedonic gLMS ranged from −79mm, indicating “strongest imaginable dislike” to 79mm, indicating “strongest imaginable like”, with 0mm indicating “neutral”. The other markers on the scale were at −42mm (“very strong dislike”), −29mm (“strong dislike”), −13mm (“moderate dislike”), −5mm (“weak dislike”), 5mm (“weak like”), 13mm (“moderate like”), 29mm (“strong like”), and 42mm (“very strong like”).

The “sensory gLMS”, measuring intensity, ranged from “no sensation” to “strongest imaginable sensation of any kind”. The sensory gLMS was scored by measuring the distance in mm between the X put on the scale by the participant and “no sensation” maker of the scale, which was scored as 0mm. The other anchors on this scale were at 5mm (“weak sensation”), 23mm (“moderate sensation”), 53mm (“strong sensation”), 80mm (“very strong sensation”) and 155mm “strongest imaginable sensation of any kind”).

### 2.3 Procedure

Ethical approval for this study was granted by the University Research Ethics Committee of the University of Manchester. Participants were asked to refrain from eating or drinking anything other than water for at least three hours before the study, in an attempt to ensure that they were experiencing comparable levels of hunger. At the start of the experiment, participants were asked to rate their hunger, fullness, craving, and thirst on a scale from one (being the least hungry, full, etc) to ten (being the most hungry, full, etc). Then participants were given careful instructions on how to use the nine-point and gLMS scales and what the order of experimental tasks would be.

The liquids were rated on the nine-point scale and on the gLMS in two separate blocks. Half the participants first rated all the liquids using the nine-point valence and intensity scales, and then repeated these ratings in the same order with the hedonic and sensory gLMS in a second block. The order of these blocks was reversed for the other half of participants. Participants always rated the pleasantness of the liquid first, and then the intensity. All stimuli were served at room temperature, in small disposable plastic cups, with ∼50 ml in each cup. Participants were asked to take a sip from the cup, and then rate the liquid on its pleasantness and intensity. We did not measure how much participants drank from each cup. Once participants had rated the liquid they were prompted to rinse their mouths with bottled water before continuing. Each stimulus was served once in each block.

The order of the stimuli was randomised, and the same randomised order was used for each participant and each block. The administration and rating of the stimuli was self-paced. Participants were given a short break in between the two blocks. After the rating of the liquids was completed participants once again rated their hunger, fullness, craving, and thirst. Then, their height and weight were measured.

### 2.4 Statistical analysis

The data were analysed in SPSS 22 (IBM Corp, 2013). Most of the data were not normally distributed (Kurtosis > 1), so non-parametric tests were used to analyse the data. One-sample Wilcoxon signed rank tests were carried out to compare the valence and intensity ratings of the five liquids to the neutral point on the hedonic scales and the least intense point on the sensory scales. To compare the three nominally-neutral liquids to each we conducted four one-way Friedman related-samples ANOVAs, a non-parametric equivalent to repeated-measured ANOVAs. Where significant, Friedman ANOVAs were followed up with related-samples Wilcoxon signed rank post-hoc comparisons. As each post-hoc analysis would involve three Wilcoxon signed rank tests, a Bonferroni correction of .017 was applied.

## 3. Results

Mean pleasantness ratings are given in Figure 2. The apple juice was rated as significantly more pleasant than neutral on both scales (both p<.001). The salty bottled water was rated as significantly more unpleasant than neutral on both scales (both p<.001). The distilled water with 2 mM NaHCO_3_ and 15 mM KCl was rated as significantly more unpleasant than neutral on the 9-point scale (p=.001), and on the hedonic gLMS (p=.041). The distilled water with 2.5 mM NaHCO_3_ and 25 mM KCl was also rated as significantly less pleasant than neutral on both the 9-point scale (p<.001), and on the hedonic gLMS (p=.001). Bottled water was not rated as significantly different from neutral on either the nine-point scale (p=.269) or on the hedonic gLMS (p=.080)^1^. In summary, participants’ ratings suggested that they felt that only the bottled water were neutral.

**Figure 2.**
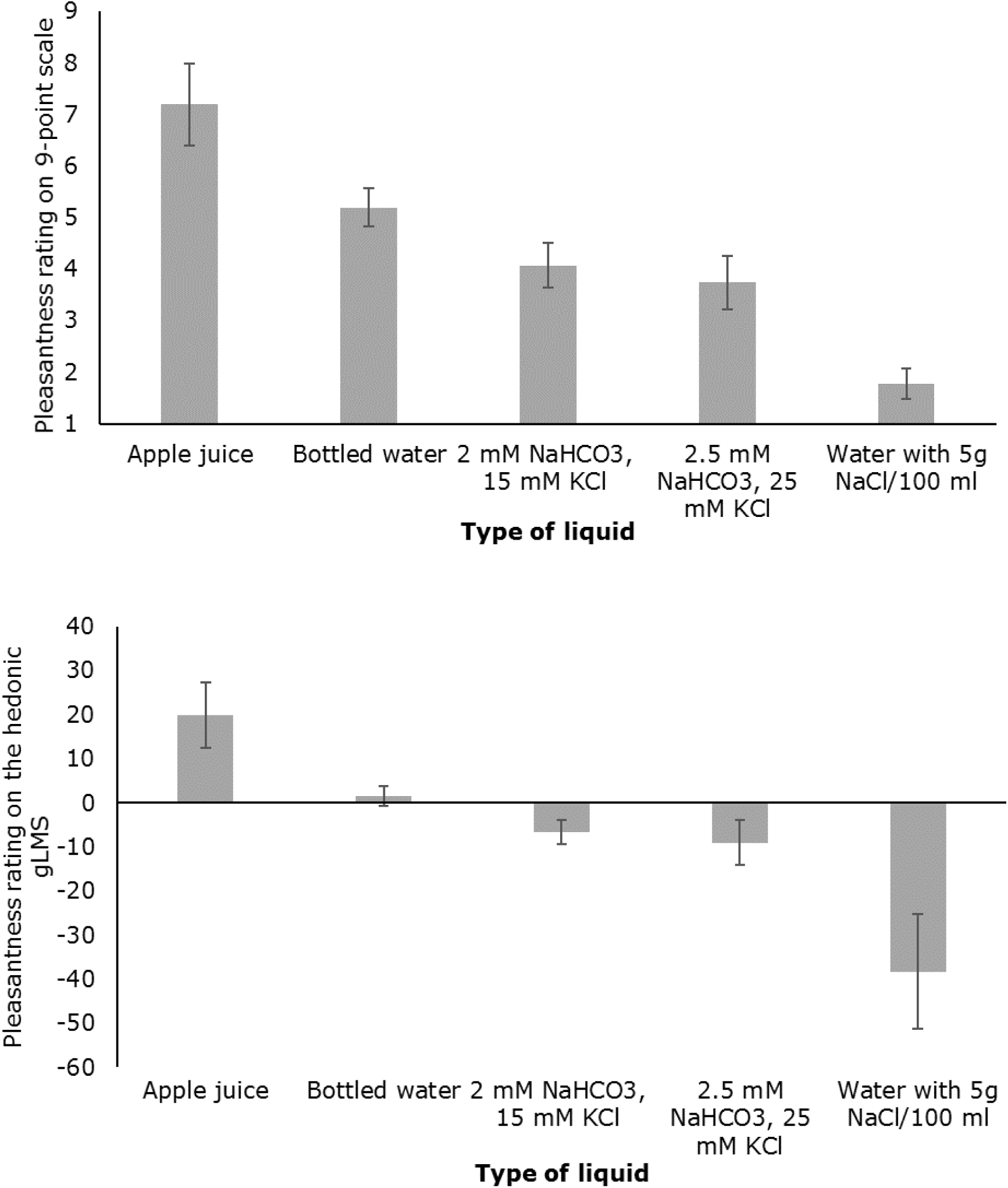
Mean pleasantness ratings for the two 9-point and gLMS scales. The graph above shows the pleasantness ratings on the 9-point scale. This scale ranged from 1 to 9, with higher ratings indicating increased taste pleasantness, and 5 being the neutral point on the scale. The graph below shows the pleasantness ratings on the hedonic gLMS. The hedonic gLMS ranged from −79 to 79, with higher ratings indicating increased taste pleasantness, and 0 being the neutral point on the scale.

Mean intensity ratings are given in Figure 3. The apple juice was rated as significantly more intense than the ‘no sensation’ mark on both scales (both p<.001). The salty bottled water was also rated as significantly more intense than ‘no sensation’ on both scales (both p<.001). The distilled water with 2 mM NaHCO_3_ and 15 mM KCl was rated as more intense than ‘no sensation’ on both scales (both p<.001). This was also the case for the distilled water with 2.5 mM NaHCO_3_ and 25 mM KCl (both p<.001), and the bottled water (p<.001 on the nine-point scale and p=.003 on the sensory gLMS). In summary, participants perceived all of the liquids to induce a distinct sensation. None of the three neutral tastes mimicked saliva’s characteristic of not being noticeable in the mouth.

**Figure 3.**
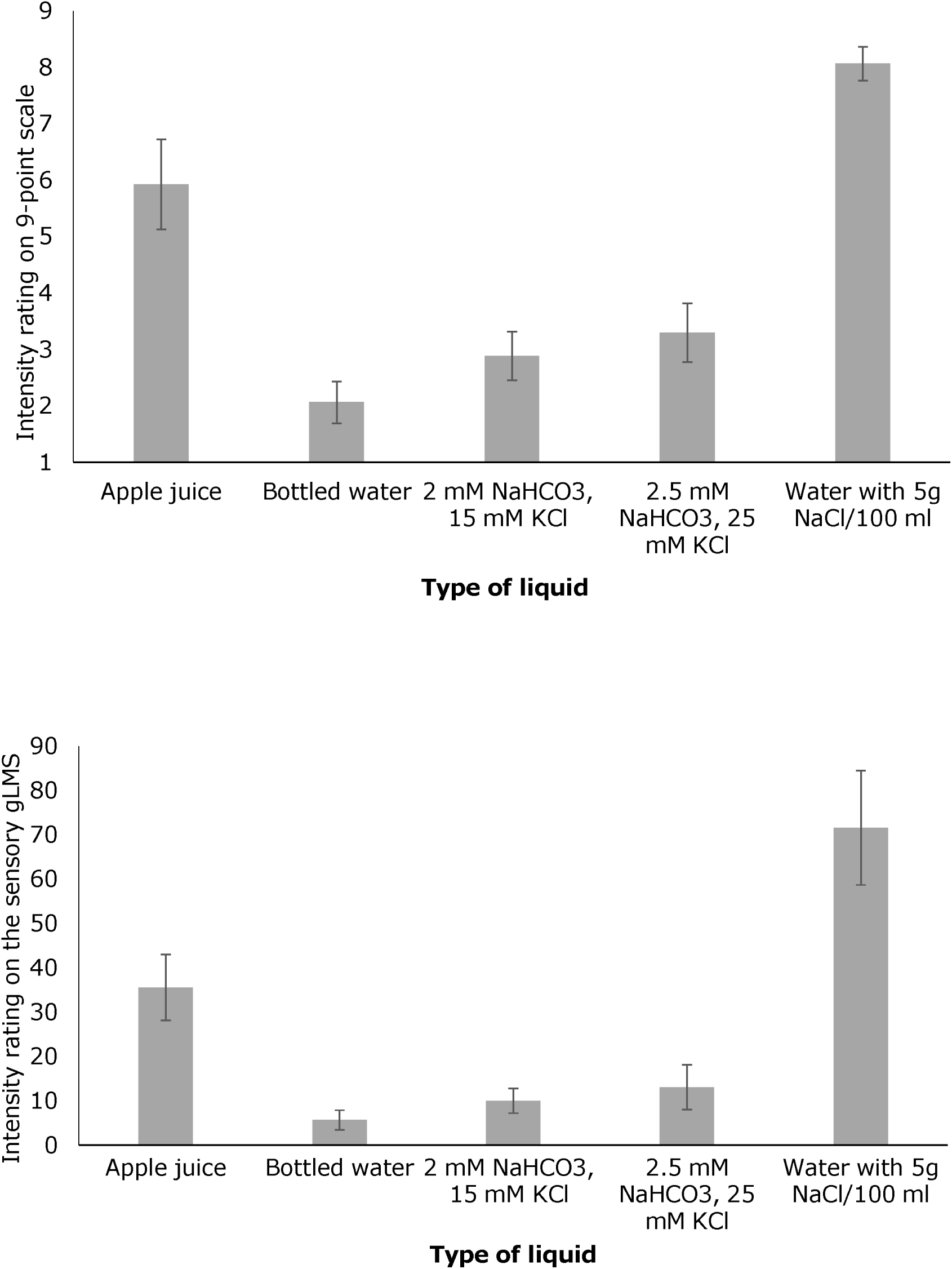
Mean taste intensity ratings. The graph above shows the intensity ratings on the 9-point intensity scale. This scale ranged from 1 to 9, with higher ratings indicating stronger taste intensity. The graph below shows the sensory gLMS, which ranged from 0 to 155, with higher ratings signifying a more intense taste.

The Friedman ANOVAs showed that the three nominally-neutral liquids differed in their pleasantness ratings on the hedonic gLMS (χ^2^ (2, N=27)=10.18, p=.006, Kendall’s W=.188) as well as their pleasantness ratings on the nine-point valence scale (χ^2^ (2, N=27)=20.69, p<.001, Kendall’s W=.383). Post-hoc tests showed that bottled water was rated as significantly more pleasant than the distilled water with 2 mM NaHCO_3_ and 15 mM KCl both on the hedonic gLMS (Z=2.50, p=.013) and the nine-point scale (Z=3.21, p=.001). Bottled water was also rated as significantly more pleasant than the distilled water with 2.5 mM NaHCO_3_ and 25 mM KCl on both the hedonic gLMS (Z=3.33, p=.001) and the nine-point valence scale (Z=3.67, p<.001). The two distilled-water solutions did not differ significantly from one another on the hedonic gLMS (Z=1.07, p=.283) or the nine-point valence scale (Z=1.47, p=.142).

The Friedman ANOVA testing the intensity ratings showed significant differences between the nominally-neutral liquids on the sensory gLMS (χ^2^ (2, N=27)=14.69, p=.001, Kendall’s W=.272) as well as the sensory nine-point scale (χ^2^ (2, N=27)=8.58, p=.014, Kendall’s W=.159). Post-hocs tests revealed that bottled water was rated as less intense than the distilled water with 2 mM NaHCO_3_ and 15 mM KCl liquids on the sensory gLMS (Z=3.33, p=.001) and the nine-point intensity scale (Z=3.21, p=.001). Bottled water was also rated as less intense than the distilled water with 2.5 mM NaHCO_3_ and 25 mM KCl on the sensory gLMS (Z=3.57, p<.001) as well as the nine-point scale (Z=3.09, =.002). There was no difference in intensity between the two distilled water solutions on the sensory gLMS (Z=1.14, p=.253) or the nine-point scale (Z=0.36, p=.719).

## 4. Discussion

This study compared the pleasantness and intensity ratings for bottled water, distilled water with 2 mM NaHCO_3_ and 15 mM KCl, and distilled water with 2.5 mM NaHCO_3_ and 25 mM KCl, on a nine-point rating scale and the hedonic and sensory gLMS. Across the board, bottled water was rated as being the most neutral of the three liquids. Distilled water with 2 mM NaHCO_3_ and 15 mM KCl was rated as mildly aversive on both scales, as was distilled water with 2.5 mM NaHCO_3_ and 25 mM. The data also showed that the bottled water was rated as the least intense-tasting liquid of the three, and that it was experienced as being significantly less intense than both distilled-water solutions on the sensory gLMS. On the nine-point intensity scale, bottled water was rated as less intense than the distilled water with 2.5 mM NaHCO_3_ and 25 mM KCl, but it did not differ in intensity from the distilled water with 2 mM NaHCO_3_ and 15 mM KCl. To the authors’ knowledge, this is the first study to have distilled water with NaHCO_3_ and KCl rated on their pleasantness and intensity. Using two separate scales, it was shown that both commonly-used concentrations of ‘artificial saliva’ were not actually rated as neutral but as mildly aversive, and that bottled water may be considered more neutral in both valence and intensity.

The fact that distilled water with NaHCO_3_ KCl and was rated as aversive is incompatible with claims that it is neutral, and casts doubt on its use as the control stimulus in experiments (e.g. Franken et al., 2011; Grabenhorst, D’Souza, et al., 2010; Grabenhorst et al., 2008; Rolls et al., 2015). In the current study, we were not able to establish why this taste was rated as aversive. It is possible that the absence of taste when consuming distilled water with NaHCO_3_ and KCl might be such an unfamiliar experience that participants perceive the experience as aversive. The fact that the more familiar bottled water was not rated as significantly aversive supports this interpretation.

It is possible the amount of liquid is an influential factor. The usual experience with bottled water is through drinking, thus people are familiar with a relatively large volume in the mouth. Saliva, in contrast, is usually present in much lower volume. The amount of liquid in the mouth in each trial of this experiment was more similar to everyday drinking, which is more liquid than is usually given to participants in gustatory research. This could emphasise the unfamiliarity of the two artificial saliva stimuli and provoke an aversive rating.

If distilled water with KCl and NaHCO_3_ is indeed rated as aversive because perceiving an absence of taste is an unfamiliar experience, experimenters who use distilled water with NaHCO_3_ and KCl as a control condition could familiarise participants with the taste prior to testing, in order for the taste to really be perceived as neutral. Another alternative for researchers could be to consider using pre-experimentally-familiar neutral liquids such as bottled water. While this liquid does not mimic the composition of saliva, our results suggest that it is hedonically neutral. Therefore, it would make a good control condition for stronger pleasant and unpleasant tastes. Alternatively, it might be worthwhile to create a liquid that mimics the properties of saliva more closely than distilled water with added NaHCO_3_ and KCl. The composition of saliva is complex, and in addition to electrolytes such as NaHCO_3_ and KCl, it also contains a range of enzymes and amino acids, among other compounds (Carpenter, 2013; Humphrey & Williamson, 2001). It might be the case that a liquid that mimics the properties of saliva more closely would be rated as hedonically neutral, and could therefore be used as a control condition in studies investigating gustatory processing.

In conclusion, this was the first study to formally test the pleasantness and intensity of two supposedly hedonically neutral tastes that are often used in gustatory research as a control condition. We found that these tastes might be experienced as sensorily meaningful, as they were perceived as intense and mildly aversive. To circumvent this issue in future gustatory research, we have recommended that researchers use still, room temperature bottled water as their control taste, or that they use a familiarisation procedure to acquaint the participants with the artificial saliva. In summary, these results suggest that using distilled water with NaHCO_3_ and KCl as a neutral control condition in gustatory research is potentially misguided.

## Footnotes

1 When analysing this data without excluding outliers (i.e. with all 30 participants), bottled water was rated as significantly more pleasant than the neutral point on the hedonic gLMS (p=.025). None of the other Wilcoxon signed-rank results changed as a result of including the outliers.

